# Dense granule protein, GRA64 interacts with host cell ESCRT proteins during *Toxoplasma gondii* infection

**DOI:** 10.1101/2021.11.02.467042

**Authors:** Joshua Mayoral, Rebekah B. Guevara, Yolanda Rivera-Cuevas, Vincent Tu, Tadakimi Tomita, Julia Romano, Leslie Gunther-Cummins, Simone Sidoli, Isabelle Coppens, Vernon B. Carruthers, Louis M. Weiss

**Author notes:** Corresponding author, Albert Einstein College of Medicine, 1300 Morris Park Avenue, Bronx, NY, 10461, USA. These authors contributed equally (Joshua Mayoral, Rebekah B. Guevara).

## Abstract

The intracellular parasite *Toxoplasma gondii* adapts to diverse host cell environments within a replicative compartment that is heavily decorated by secreted proteins. In attempts to identify novel parasite secreted proteins that influence host cell activity, we identified and characterized a trans-membrane dense granule protein dubbed GRA64 (TGME49_202620). We found that GRA64 is on the parasitophorous vacuolar membrane (PVM) and is partially exposed to the host cell cytoplasm in both tachyzoite and bradyzoite parasitophorous vacuoles. Using co-immunoprecipitation and proximity-based biotinylation approaches, we demonstrate that GRA64 appears to interact with certain components of the host Endosomal Sorting Complexes Required for Transport (ESCRT). Genetic disruption of GRA64 does not affect acute *Toxoplasma* virulence in mice nor encystation as observed via tissue cyst burdens in mice during chronic infection. However, ultrastructural analysis of Δ*gra64* tissue cysts using electron tomography revealed enlarged vesicular structures underneath the cyst membrane, suggesting a role for GRA64 in organizing the recruitment of ESCRT proteins and subsequent intracystic vesicle formation. This study uncovers a novel host-parasite interaction that contributes to an emerging paradigm in which specific host ESCRT proteins are recruited to the limiting membranes (PVMs) of tachyzoite and bradyzoite vacuoles formed during acute and chronic *Toxoplasma* infection.

**IMPORTANCE:** *Toxoplasma gondii* is a widespread foodborne parasite that causes congenital disease and life-threatening complications in immune compromised individuals. Part of this parasite’s success lies in its ability to infect diverse organisms and host cells, as well as to persist as a latent infection within parasite constructed structures called tissue cysts. In this study, we characterized a protein secreted by *T. gondii* into its parasitophorous vacuole during intracellular infection, which we dub GRA64. On the vacuole, this protein is exposed to the host cell and interacts with specific host ESCRT proteins. Parasites lacking the GRA64 protein exhibit ultrastructural changes in tissue cysts during chronic infection. This study lays the foundation for future studies on the mechanics and consequences of host ESCRT-parasite protein interactions.

## INTRODUCTION

The intracellular parasite *Toxoplasma gondii* is a widespread pathogen estimated to infect approximately one-third of human beings worldwide (1). The extensive prevalence of *T. gondii* can be partially attributed to its flexible life cycle, one in which a wide variety of hosts can be infected most commonly by the ingestion of material containing parasites within cystic structures (2, 3). Within intermediate hosts (where asexual replication occurs), *T. gondii* interconverts between two life stages that typify acute and chronic infection (4). The tachyzoite life stage spreads throughout the body of the host during acute infection and causes disease through the lysis of host cells and subsequent tissue damage, whereas the bradyzoite life-stage exhibits a quasi-dormant existence within intracellular tissue cysts predominately in the brain and muscle tissue during chronic infection (5). Bradyzoite differentiation is induced by various factors that can be summarized as stressors to the parasite which trigger a bradyzoite transcriptional program and an altered pattern of intracellular replication (6, 7). The absence of stressors, as might occur in the setting of compromised host immunity, is a permissive state that allows for bradyzoite re-conversion back to the tachyzoite stage, leading to life-threatening encephalitis and other complications (8, 9). As there are currently no available treatments that eradicate tissue cysts from chronically infected hosts, there is a major need to understand the processes that contribute to parasite latency.

Both tachyzoite and bradyzoite life stages replicate within specialized intracellular vacuoles termed either the parasitophorous vacuole (during tachyzoite infection) or the tissue cyst (during bradyzoite infection) (4). Parasitophorous vacuoles and tissue cysts are extensively modified by the parasites within these compartments through the secretion of lipids and proteins into the vacuolar space and cyst matrix (10, 11). For example, the secretion of multi-lamellated vesicles from the basal end of tachyzoites gives rise to the intravacuolar network (IVN) (12), which is known to play a pivotal role in the acquisition of host cell resources (13, 14). A seemingly analogous structure, the intracystic network, has been described in tissue cysts as well (11). The defining feature of tissue cysts historically has been the cyst wall, which appears as an electron dense conglomeration of vesicular and filamentous material that underlies the cyst membrane (11). Despite the ultrastructural characterization of the tachyzoite parasitophorous vacuole and bradyzoite tissue cysts, much remains to be discovered regarding the number and function of the proteins secreted into these specialized vacuoles and their contribution to parasitism. Many of the proteins found either in a soluble or insoluble, membrane-associated state arise from secretory organelles termed dense granules (15), which constitutively release “GRA” proteins throughout intracellular development (16). Many GRA proteins do not contain domains of known function, suggesting that novel processes unique to the parasite’s intracellular niche might be mediated by these proteins.

Despite their enigmatic nature, the function of very few GRA proteins have been elucidated in vesicle trafficking and nutrient acquisition. GRA2 and GRA6 are pivotal in forming the IVN during the early stages of tachyzoite parasitophorous vacuole development (17). GRA7 sequesters host endocytic organelles to the parasitophorous vacuole membrane (PVM) (18). GRA3 recruits host Golgi and aids in the trafficking of Golgi vesicles across the PVM (19), and MAF1 recruits and tethers host mitochondria to the PVM (20). GRA17 and GRA23 have been shown to traffic small molecules across the PVM (21), and it is suspected that GRA17 serves the same purpose during bradyzoite development (22). Recently discovered is the interaction between GRA14 and the host Endosomal Sorting Complex Required for Transport (ESCRT) machinery, which mediates ESCRT-dependent virus-like particle budding and internalization of host cytosolic proteins at the PVM (23). Certain GRA proteins have been shown to be partially exposed to the host cell following their transmembrane insertion into the PVM, such as GRA5 (24), GRA6 (25), and GRA14 (26). GRA6 has been shown to influence host cell NFAT activity (27), while a specific GRA15 allele has been shown to induce host cell NF-κB activity (28), presumably through host cell exposure of GRA15 at the PVM. Although GRA15 is the main regulator of NF-κB, GRA7 and GRA14 also modify nuclear localization of RelA/p65, a member of the NF-κB, complex, from the PVM (29). Certain GRA proteins have even been found to operate beyond the PVM interface, entering the host cell cytoplasm and nucleus and influencing host cell function by interacting with specific host cell proteins (30, 31).

In efforts to identify novel secreted *Toxoplasma* proteins that may directly influence host cell activity, we devised an *in silico* screen to predict genes encoding proteins with properties similar to known exported effector proteins. We identified one protein from this screen secreted into the vacuole, but not beyond the PVM or cyst membrane, likely due to the presence of a transmembrane domain. We set about characterizing this novel protein, TGME49_202620 (dubbed GRA64), in detail and found that its N-terminus is exposed to the host cell during intracellular infection. Co-immunoprecipitation and proximity-based biotinylation approaches revealed that the most frequent host cell interacting partners of GRA64 are components of the <u>E</u>ndosomal <u>S</u>orting <u>C</u>omplexes <u>R</u>equired for <u>T</u>ransport (ESCRT), which canonically generate intraluminal vesicles away from the cytosol following their stepwise recruitment. Genetic deletion of GRA64 did not impair tachyzoite growth, nor did it impact cyst burden during chronic infection. Ultrastructural analysis, however, of Δ*gra64* tissue cysts demonstrated enlarged cyst membrane adjacent intraluminal vesicles compared to wild type tissue cysts. Electron tomography of Δ*gra64* tissue cysts revealed these enlarged intraluminal vesicles to be cyst membrane invaginations, suggesting perturbed ESCRT-mediated scission events at the cyst membrane in the absence of GRA64. However, unlike GRA14 (23), GRA64 neither participates in ESCRT-dependent virus like particle budding nor regulates ingestion of host cytosolic proteins. Altogether, our findings suggest that GRA64 is one of several membrane bound GRA proteins facing the host cell that recruits host ESCRT, the consequences of which have yet to be fully understood.

## RESULTS

### Discovery of a novel GRA protein, GRA64

The gene TGME49_202620 was identified from an *in silico* screen aimed at characterizing novel exported parasite effector proteins. The gene is predicted to encode a protein containing a signal peptide and a transmembrane domain proximal to the C-terminus (Fig. 1A). As assessed by the webserver IUPred3 (32), the gene product is predicted to contain regions of protein intrinsic disorder (Fig. 1B), suggesting the capability of promiscuous protein-protein interactions. The low disordered regions correspond to the signal peptide and the transmembrane domain of TGME49_202620 (Fig. 1B). Endogenously tagged PruQ parasites were engineered with a 3xHA tag appended to the C-terminus of TGME49_202620, upstream of the stop codon. Immunofluorescence assays (IFAs) of C-terminus tagged TGME49_202620 protein demonstrate secretion into the lumen of the parasitophorous vacuole, where it was frequently found to outline the parasitophorous vacuole membrane (PVM) under tachyzoite growth conditions using Type II PruΔ*ku80*Δ*hxgprt* (PruQ) parasites (Fig. 1C, top panel). Endogenously tagged PruQ parasites were engineered with a 3xHA tag appended to the N-terminus of TGME49_202620, downstream of the predicted signal peptide. IFAs of extracellular parasites from this strain revealed that the TGME49_202620 protein co-localizes with the dense granule marker GRA1, indicating that this protein is likely packaged into dense granules prior to secretion into the parasitophorous vacuole (Fig. 1C, bottom panel). Given this result, we hereafter refer to the TGME49_202620 gene product as GRA64. To assess the localization of GRA64 more accurately during intracellular infection, immunoelectron microscopy of tachyzoite vacuoles was performed. The images demonstrate that GRA64 signal is most frequently detected in association with membranous tubular structures reminiscent of the intravacuolar network (IVN) (Fig. 1D). No PVM labeling was observed in this experiment, although the integrity of the PVM appeared to be compromised during the preparation of these samples likely due to the use of Triton X-100. To determine whether the predicted transmembrane domain conferred membrane interacting properties to GRA64, fractionation experiments were performed using material from infected monolayers containing tachyzoites. GRA64 was found to predominately associate with the high-speed pellet fraction, with only trace amounts present in the high-speed supernatant (Fig. 1E). Treatment with 6M Urea, 1M NaCl, 0.1M Na_2_CO_2_, 0.1% NP-40, and 0.1% Triton X-100 revealed that only 6M urea and non-ionic detergents (NP-40, Triton X-100) were capable of dissociating modest amounts of GRA64 protein from the high-speed pellet fraction, indicating that GRA64 exhibits integral membrane protein properties (Fig. 1E), similar to what has been described for other GRA proteins with transmembrane domains such as GRA5 (24). The immunoblot shown in Fig.1E demonstrates that GRA64 typically migrates slightly below 55kDa, slightly slower than the predicted size of ∼39kDa. GRA64 has a predicted *N*-glycosylation site at amino acid 55 and several phosphorylation sites detected in prior proteomic datasets deposited onto the *Toxoplasma* database ToxoDB (33), and this could account for slower migration together with the intrinsically disordered structure predicted by IUPred3. No differences were observed in GRA64 migration from protein harvested from extracellular parasites, a 24-hour infected tachyzoite culture, or a 3-day induced bradyzoite culture (data not shown).

**Figure 1.**
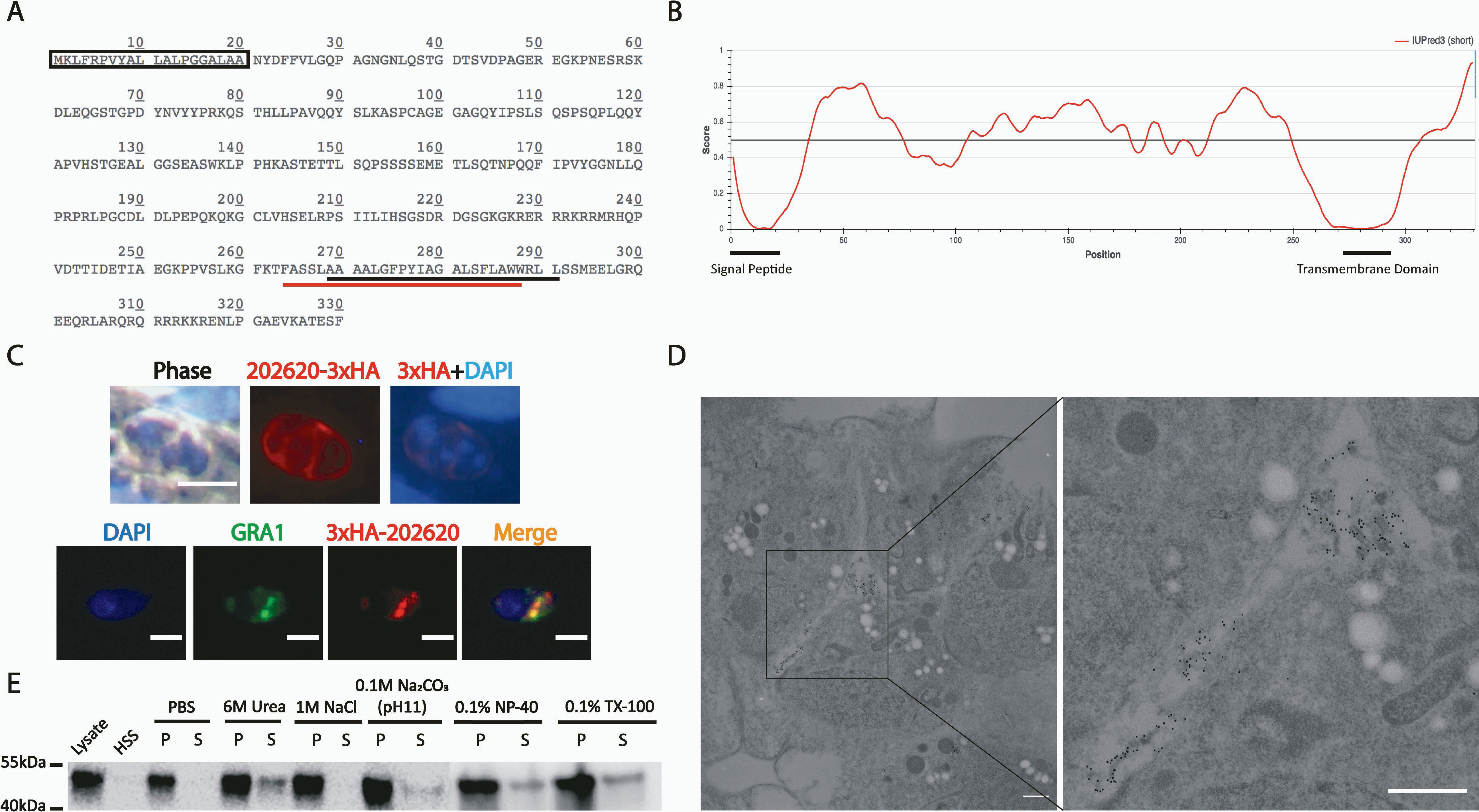
GRA64 is a dense granule integral membrane protein localizing to the intravacuolar network during tachyzoite infection. **(A)** Amino acid sequence of TGME49_202620. The predicted signal peptide is indicated by a box (SignalP 5.0 prediction), while the predicted transmembrane domains are indicated by underlines (black – TMHMM prediction, red – Phobius prediction). **(B)** Results from IUPred3 short disorder prediction of TGME49_202620 protein intrinsic disorder is displayed using medium smoothing. The score indicates regions of low and high disorder. **(C)** Top panel - IFA image of C-terminus 3xHA epitope tagged TGME49_202620 protein from tachyzoite vacuoles 1-day post-infection. TGME49_202620 is detected within the parasitophorous vacuole, seemingly outlining the PVM. Bars, 10µm. Bottom panel – IFA image of extracellular parasites expressing N-terminus 3xHA epitope tagged TGME49_202620 protein. TGME49_202620 (hereafter referred to as GRA64) colocalizes with GRA1. Bars, 5µm. **(D)** Immunoelectron microscopy of a tachyzoite vacuole expressing 3xHA epitope tagged GRA64 protein. Signal from 10nm gold particles conjugated to anti-HA antibody appear to associate with membranous structures of the IVN within the parasitophorous vacuole. Bars, 500nm. **(E)** Immunoblot of GRA64 protein detected in various fractions following ultracentrifugation of protein lysates from tachyzoite infected fibroblast monolayers. HSS or S – high speed supernatant; P – high speed pellet (membrane fraction). Treatment of high-speed pellets with 6M Urea, 1M NaCl, 0.1M Na_2_CO_3_ (pH 11), 0.1% NP-40, and 0.1% Triton X-100 (TX-100).

### N-terminus of GRA64 is exposed to the host cell cytoplasm during intracellular infection

We hypothesized that GRA64 is exposed to the host cell cytoplasm following insertion of the transmembrane domain into the lipid membranes of the PVM or IVN. Parasites expressing an N-terminal HA tagged version of GRA64 at the endogenous locus were used in selective permeabilization IFA experiments (Fig. 2A schematic). At the digitonin concentration of 0.001% (w/v) to selectively permeabilize the host cell plasma membrane but not the PVM to antibodies, GRA64 signal was detected in intact parasitophorous vacuoles, as assessed by the lack of labeling of MAG1, an abundant vacuolar protein (Fig. 2A, Digitonin Perm panels). GRA64 exposure to the host cell cytoplasm was detected both under tachyzoite (pH7) and bradyzoite (pH8) growth conditions (Fig. 2A).

**Figure 2.**
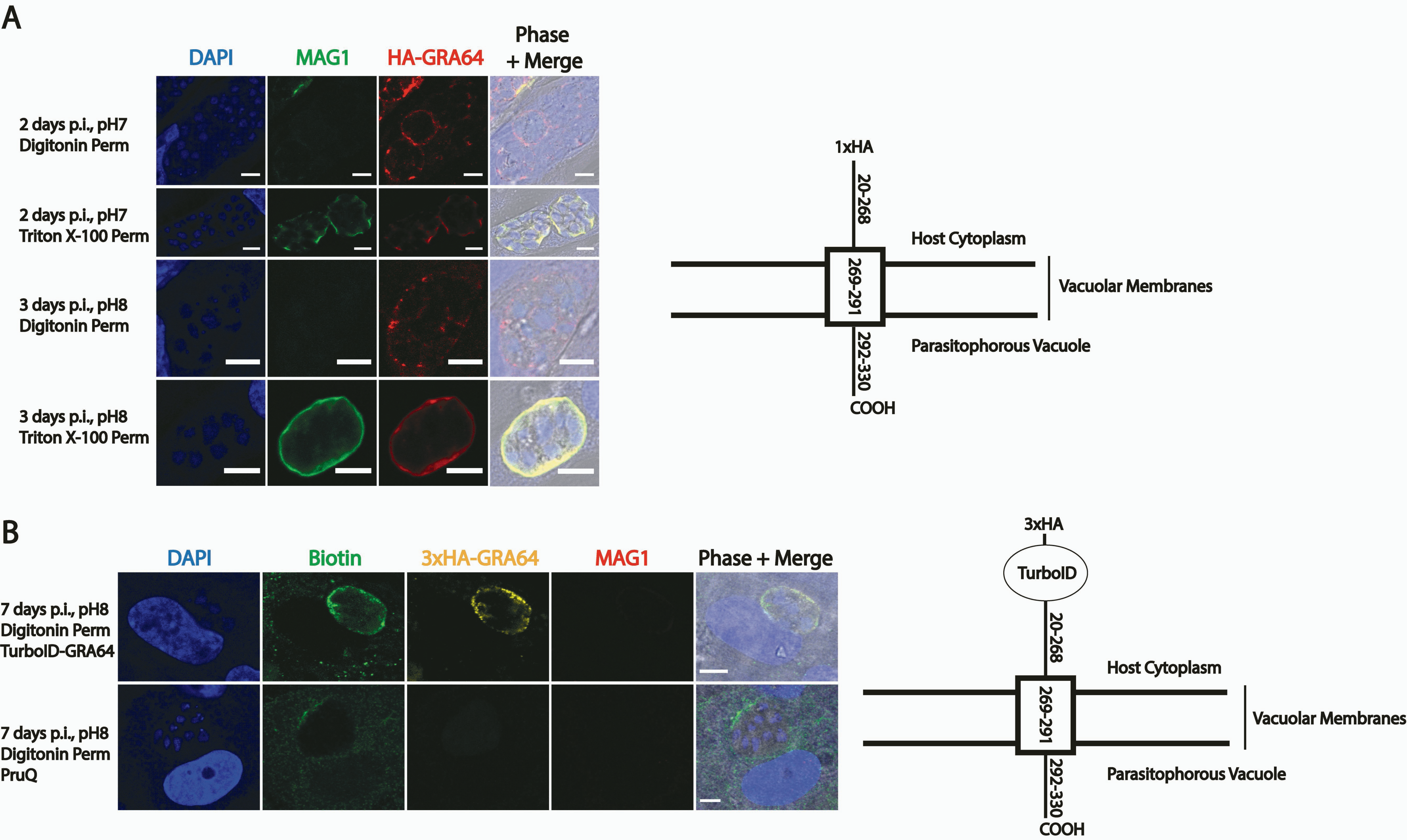
Epitope tagged- and TurboID tagged-GRA64 proteins are exposed to the host cell cytoplasm during intracellular infection. **(A)** IFA panels of parasites expressing GRA64 tagged at the N-terminus with a 1xHA tag in tachyzoite vacuoles (2 days p.i., pH7) or induced bradyzoite vacuoles (3 days p.i., pH8). The digitonin concentration used for permeabilization in these experiments (0.001%) selectively permeabilizes the host cell membrane, but not the PVM. In contrast, Triton X-100 fully permeabilizes the host cell and the PVM, as demonstrated by positive staining of MAG1, a parasite protein localizing to the lumen of parasitophorous vacuoles and to the cyst wall in differentiating parasite vacuoles. Bars, 5µm. **(B)** IFA panels of parasites expressing GRA64 tagged at the N-terminus with a 3xHA and the proximity-based biotinylating enzyme TurboID tag or untagged PruQ parasites in induced bradyzoite vacuoles (7 days p.i., pH8). Infected cells were permeabilized with digitonin (0.001%) to selectively permeabilizes the host cell membrane, but not the PVM. Vacuoles were stained with αbiotin and αHA to confirm activity of the TurboID tag and expression of GRA64 via the 3xHA tag at the host cytoplasm interface, respectively. The lack of MAG1 labeling was used as an indicator for selective permeabilization. Bars, 5µm.

With the aim of eventually identifying host cell proteins that potentially interact with GRA64, parasites expressing a TurboID-GRA64 fusion protein were engineered. The proximity-based biotinylating enzyme TurboID (34) and a 3xHA epitope tag were appended to the N-terminus of endogenous GRA64 (Fig. 2B, schematic). To assess whether the addition of the TurboID tag interfered with the topology of the GRA64 protein, selective permeabilization experiments were again performed with this parasite strain under bradyzoite growth conditions supplemented with exogenous biotin. TurboID-GRA64 fusion protein was also detected in intact parasitophorous vacuoles, as determined by the detection of 3xHA signal in vacuoles without MAG1 signal (Fig. 2B). Biotinylation was also detected in intact parasitophorous vacuoles using anti-biotin antibody, indicating that the TurboID enzyme is also active specifically at the host cell exposed vacuolar membrane interface. Hence, the N-terminal portion (with respect to the predicted transmembrane domain) of various GRA64 fusion proteins appears to be consistently exposed to the host cell cytoplasm.

### GRA64 interacts with host cell proteins from the Endosomal Sorting Complexes Required for Transport (ESCRT)

Given the host cell exposure of GRA64, we hypothesized that certain host cell proteins may interact with the GRA64 protein. Co-immunoprecipitations (Co-IPs) were performed using protein lysates from tachyzoite-infected human fibroblast cultures, using PruQ parasites expressing endogenously tagged 3xHA-GRA64 protein (24 hours post-infection). Co-IPs of GRA64 in protein lysates from bradyzoite-infected human fibroblasts (four days post-infection) and from mouse primary cortical neuron infected cultures (two days post-infection) were also performed, to determine if any host cell protein associations were common between life stages and host cell type, as well as between different host species. Untagged PruQ parasites cultured under the same conditions were used as a negative control for all experiments to identify proteins that non-specifically bound to the anti-HA antibody coated magnetic beads. Two independent experiments were performed for each condition. Following overnight incubation of harvested proteins with anti-HA magnetic beads, protein was washed, eluted with Laemmli buffer, removed from detergent and digested into peptides in S-TRAP columns, and analyzed by LC-MS/MS.

Among the parasite proteins significantly enriched from tachyzoite infected fibroblast samples (log_2_ fold-enrichment ≥ 1.0, p-value ≤ 0.10 in two independent experiments), various GRA proteins were identified, as expected based on the localization of GRA64 in the vacuole and presence in dense granules within the parasite (Fig. 1C). The only host cell proteins significantly enriched from tachyzoite infected fibroblast samples were various proteins belonging to or associated with the endosomal sorting complexes required for transport (ESCRT) (Fig. 3). Specifically, representatives from the ESCRT-I (TSG101, VPS37A, VPS28) and ESCRT-III (CHMP4B) complex were enriched in most Co-IP experiments (Fig. 3A-C), as well as proteins associated with ESCRT recruitment (PDCD6 and UMAD1). However, it is worth noting that GRA64 was not detected as significantly enriched in the neuron Co-IPs (Fig. 3C), likely due to low protein enrichment overall from neuron infected cultures, resulting in greater variability in the LC-MS/MS analysis pipeline. Despite the absence of significant bait-protein enrichment in the neuron experiment, the data demonstrate that similar parasite and host cell proteins to those found from fibroblast cultures are significantly enriched (GRA proteins and ESCRT proteins), indicating an intriguing common recruitment of host proteins at the IVN/PVM interface (and analogous structures in tissue cyst) across each condition tested.

**Figure 3.**
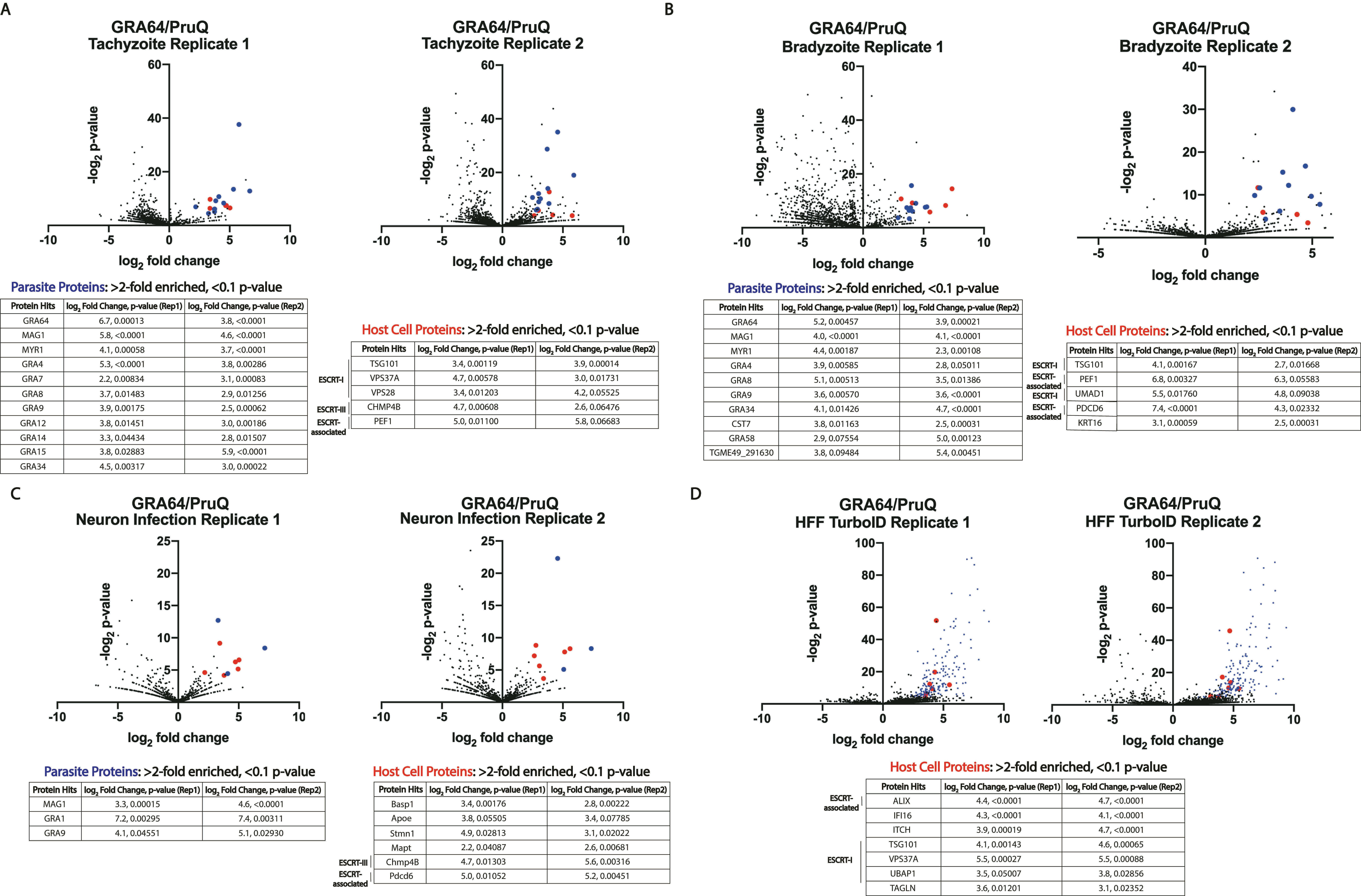
GRA64 interacts with host ESCRT proteins. Volcano plots of LC-MS/MS results from co-immunoprecipitations of endogenously tagged 3xHA-GRA64 protein from either infected human fibroblast monolayers under **(A)** tachyzoite or **(B)** bradyzoite growth conditions (1- and 4-days post-infection, respectively), or from **(C)** mouse primary cortical neuron cultures (2-days post-infection). In each volcano plot, red and blue data points indicate a host or parasite protein classified as significant in two independent experiments respectively (log_2_ fold-enrichment ≥ 2 and p-value of < 0.1 in both replicate 1 and replicate 2). Under each condition, the host proteins that were identified as significant hits were largely ESCRT or ESCRT-associated proteins. **(D)** TurboID-GRA64 expressing parasites were used to identify proximal proteins in human foreskin fibroblast cultures. A list of all host cell proteins (red dots in the volcano plot) with log_2_ fold-enrichment ≥ 2 and a p-value of ≤ 0.1 are listed in the table. Significantly enriched parasite protein hits (blue dots) are provided in Supplementary Dataset 2.

The Co-IP approach necessitates the lysis of host cells to immunoprecipitate GRA64 protein from the parasitophorous vacuole, which may lead to artificial interactions that do not normally occur during infection. To determine if ESCRT protein proximity to GRA64 could be detected in living cells, we utilized proximity-based biotinylation with TurboID, using the afore-mentioned TurboID-GRA64-expressing parasite strain (Fig. 2B) and an untagged parasite strain as a control. Streptavidin resin was used to enrich biotinylated proteins in two independent experiments, harvesting proteins from bradyzoite-induced cultures with exogenous biotin three days post-infection. The results demonstrated that three ESCRT-I proteins (TSG101, VPS37A, UBAP1) and the accessory ESCRT protein ALIX were identified as significantly enriched (log_2_ fold-enrichment ≥ 1.5, p-value ≤ 0.1) in TurboID-GRA64 cultures compared to untagged cultures grown under similar conditions in both independent experiments (Fig. 3D). Hence, both Co-IP and TurboID approaches provide evidence for an association between GRA64 and host ESCRT proteins. TurboID labeling experiments were also performed during infection in mouse primary cortical neurons. No host proteins were found to be significantly enriched in TurboID labeled samples in two independent experiments with neurons, despite evidence for successful TurboID labeling based on the significant enrichment of parasite proteins (Fig. S1). Host proteins may have been less efficiently labeled during neuron infection due to relatively lower amounts of host proteins associated with vacuolar membranes compared to the fibroblast infection model.

To help validate the results obtained by LC-MS/MS, 3xHA-GRA64 pulldowns with anti-HA beads were repeated under tachyzoite growth conditions in human foreskin fibroblast cultures and during neuron infection (Fig. 4A). A clear enrichment of TSG101 and PDCD6 were observed in the human foreskin fibroblast infection eluates from 3xHA-GRA64 samples, but not untagged control eluates (Fig. 4A, left panel). Similarly, PDCD6 was found to be enriched by GRA64 pulldown during neuron infection (Fig. 4A, right panel), confirming LC-MS/MS findings. A reciprocal Co-IP was also performed using antibody to the ESCRT-III protein CHMP4B, conjugated to magnetic dynabeads. As a control, unconjugated dynabeads were used in parallel. Incubating control and CHMP4B conjugated magnetic beads with tachyzoite infected cultures expressing 3xHA-tagged GRA64 protein demonstrated CHMP4B enrichment of GRA64 over control beads after several wash steps and elution, as determined by immunoblotting (Fig. 4B). These data suggest that the interaction between GRA64 and CHMP4B is reproducible, at least during the artificial Co-IP cell lysis and wash conditions used in this experiment. Altogether, LC-MS/MS results and pulldowns with GRA64 and CHMP4B demonstrate that GRA64 interacts with specific ESCRT-I, ESCRT-III, and accessory ESCRT proteins.

**Figure 4.**
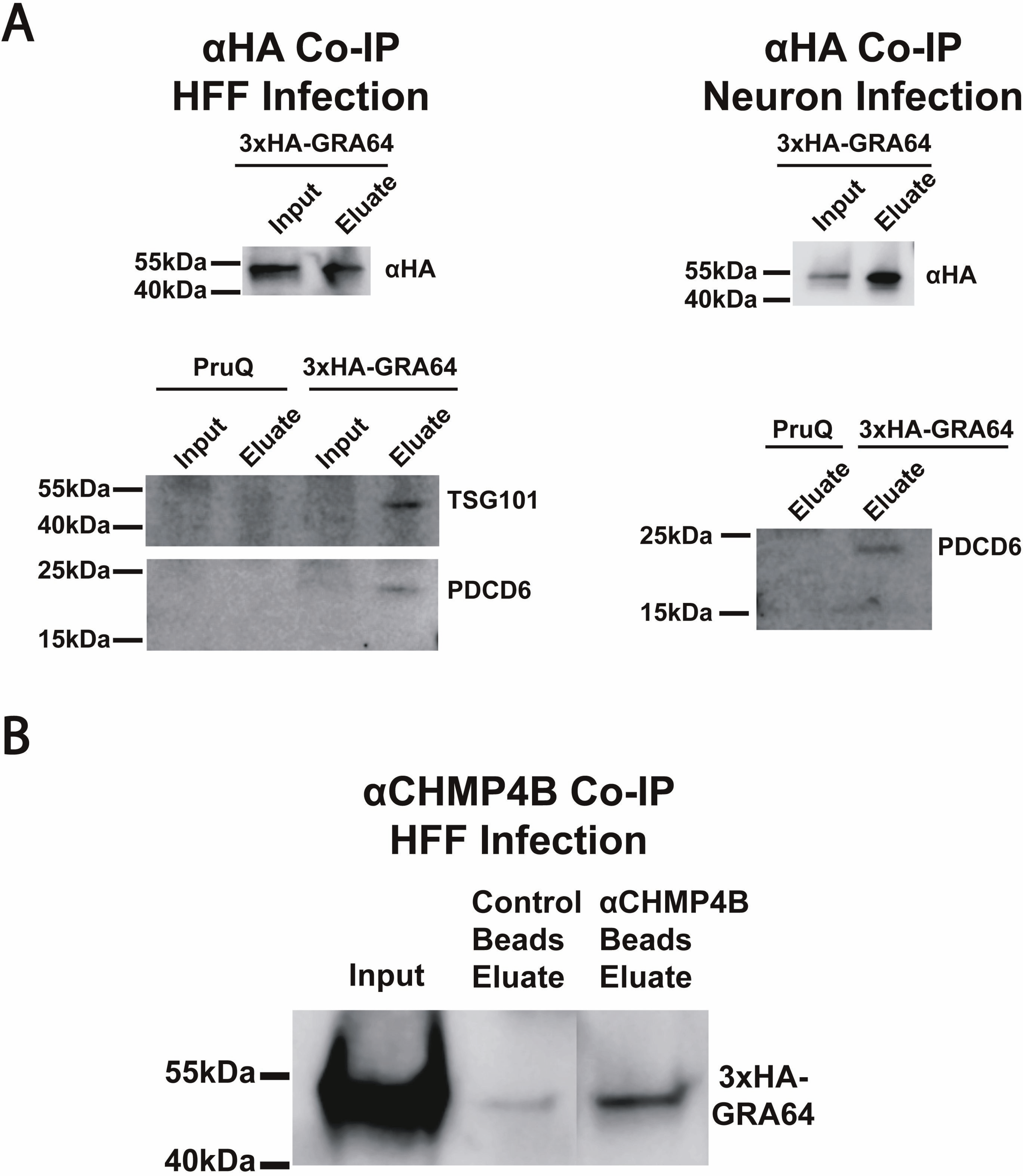
Immunoblot confirmation of ESCRT protein enrichment by GRA64. **(A)** Immunoblots of samples from an anti-HA Co-IP using either 3xHA-GRA64 tagged or untagged PruQ parasites as a control. Protein lysates were harvested for Co-IP from either human foreskin fibroblast infected monolayers (left panel) or from neuron infected cultures (right panel). Although no robust enrichment of GRA64 is seen in the eluate fraction compared to the input fraction (top left panel), both TSG101 and PDCD6 were only detected in the 3xHA-GRA64 eluate samples and not the control samples (bottom left panel), indicating ESCRT protein enrichment in agreement with LC-MS/MS results. Similarly, PDCD6 was only detected the 3xHA-GRA64 eluate and not the control eluate from neuron infection samples (bottom right panel). **(B)** Immunoblot of 3xHA-GRA64 protein immunoprecipitated with dynabeads conjugated with anti-CHMP4B antibody (and unconjugated beads, as a control). There is a notable enrichment of GRA64 using CHMP4B conjugated beads compared to unconjugated beads.

### Genetic disruption of GRA64 does not affect tachyzoite growth *in vitro* and in mice nor cyst burden in mice during chronic infection

To determine how GRA64 might contribute to parasite fitness, we next disrupted and complemented the GRA64 gene in PruQ parasites using a previously described Cas9 strategy (35). Immunoblots of protein lysates from each strain revealed, as expected, the absence of 3xHA-GRA64 protein in the Δ*gra64* and comparable amounts of GRA64 protein in the parental (3xHA-GRA64) and complemented (GRA64-COMP) strains (Fig. S2A). Neither *in vitro* growth in fibroblasts (determined by plaque assays, Fig. S2B) nor acute virulence *in vivo* (Fig. S2C) was affected in PruQΔ*gra64* parasites, indicating that GRA64 is dispensable for tachyzoite growth. We next assessed cyst burdens in CBA/J mice chronically infected with PruQ strain parasites 30 days post-infection. In two independent experiments, we observed a trend for reduced cyst burdens in mouse brains infected with the Δ*gra64* PruQ strain, compared to the parental and complemented counterparts (Fig. S3A). To further evaluate the cyst phenotype, in two independent experiments we quantified cyst burden in equivalent GRA64 strains generated in the relatively more cystogenic ME49Δ*ku80*Δ*hxgprt* (ME49Q) background. The results showed an overall increase in cyst yields, but no difference in ME49QΔ*gra64* cyst burdens compared to the complemented strain. Altogether, the data suggest GRA64 does not significantly influence cyst burden in a predictable manner (Fig. S3B).

### Tissue cysts formed by Δ*gra64* parasites demonstrate changes in cyst ultrastructure

We next evaluated the localization of GRA64 in tissue cysts formed *in vivo*. Immunogold labeling of complemented GRA64 cysts expressing 3xHA-GRA64 revealed labeling of the cyst wall region and parasite dense granules using anti-HA antibody, providing evidence that GRA64 is indeed expressed by mature bradyzoites *in vivo* (Fig. 5A). Furthermore, we assessed whether tissue cysts formed *in vivo* by Δ*gra64* parasites exhibited any morphological defects related to GRA64-ESCRT interaction. We hypothesized that defects in the recruitment of host ESCRT proteins at the cyst membrane interface could result in more prominent “stalled” cyst membrane invaginations, due to the lack of efficient ESCRT-mediated membrane scission. Using the wild-type (ME49Q), GRA64 parental (3xHA-GRA64), GRA64 knockout (Δ*gra64*), and complemented (GRA64-COMP) ME49Q strains, we harvested and purified cysts by Percoll from mouse brains chronically infected for four weeks. Post-purification and fixation, cysts were analyzed by electron microscopy. The images revealed that while wildtype, parental, and complemented cysts exhibited standard cyst wall architecture typified by electron dense material and occasionally small vesicular material underneath the cyst membrane, however Δ*gra64* cysts more frequently harbored large vesicular structures (200-400nm) proximal to the cyst membrane (Fig. 5B). To further investigate the large vesicular structures, we performed electron tomogram analysis of thicker sections from *in vivo* brain cysts (250nm) of ME49Q (Movie 1) and Δ*gra64* (Movie 2). The ME49Q tomogram demonstrates narrow cyst membrane invaginations with occasional vesicles trapped within the lumen of these invaginations. We further traced the vesicular structures in the Δ*gra64* tomogram to see if they were continuous with the cyst membrane and observed a few close vesicular-cyst membrane contacts (Fig. S4). These large vesicular structures potentially represent enlarged invaginations that have not been efficiently excised from the cyst membrane due to perturbed ESCRT recruitment. However, we cannot conclude with certainty the nature of these seemingly aberrant structures from static images alone.

**Figure 5.**
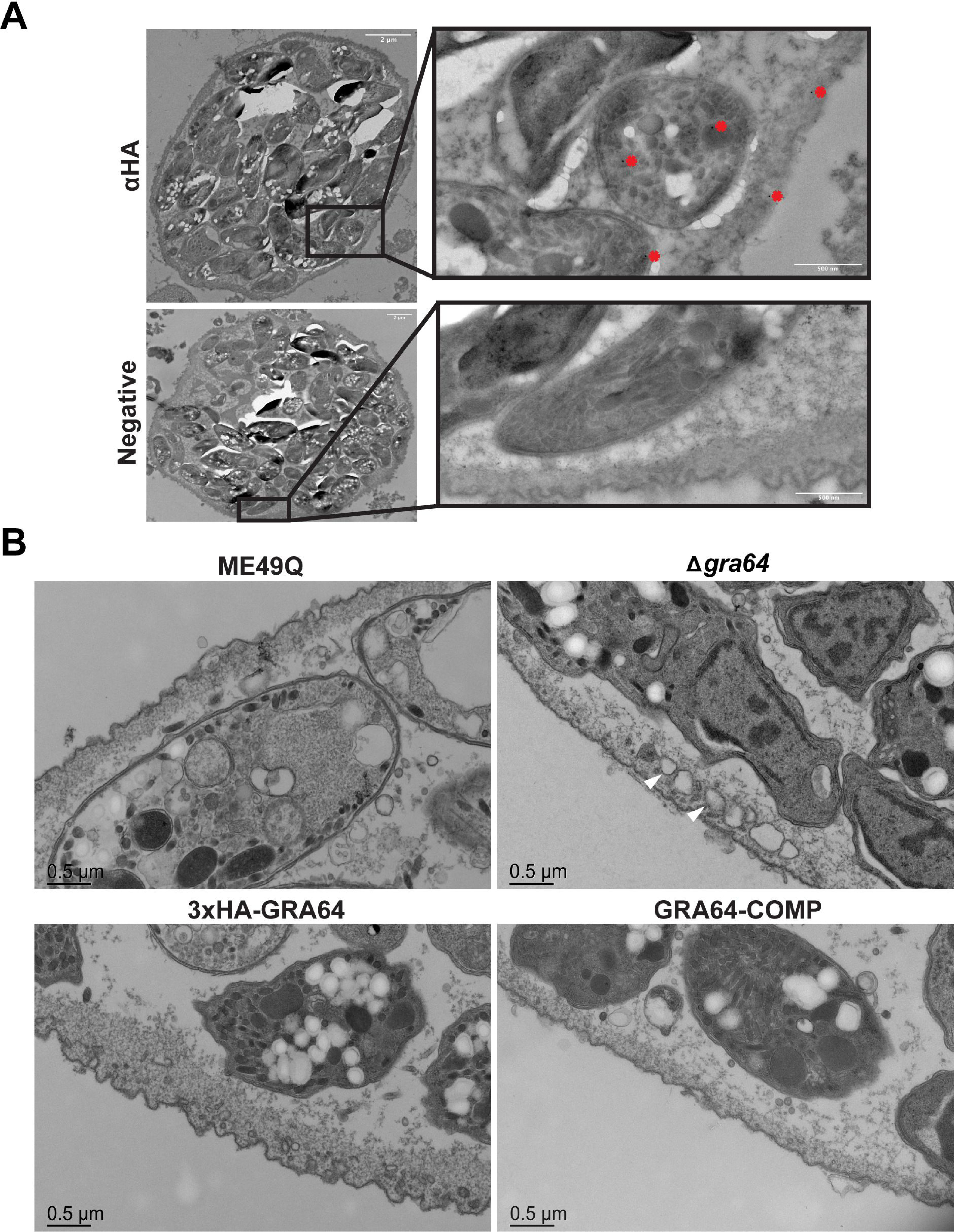
Δ*gra64* derived tissue cysts exhibit a change in ultrastructure. **(A)** Immunogold labeling (using rat anti-HA and anti-rat 10nm gold particle conjugated antibodies) of GRA64-COMP strain tissue cysts demonstrates GRA64-positive signal (red asterisks) seemingly within dense granule structures in bradyzoites and within the cyst wall. A GRA64-COMP tissue cyst labeled with only anti-rat gold conjugated antibody is provided in the panel below for comparison (“negative”). Bars = 2um and 500nm for magnified inserts. **(B)** Magnified views of tissue cyst walls and cyst membranes formed by each strain as indicated. Note the presence of large vesicular structures within the cyst and proximal to the cyst wall in the Δ*gra64* panel (white arrowheads). Bars = 0.5um.

### GRA64 does not recruit ESCRT machinery in virus-like particle assay and is not required for internalization of host cytosolic proteins

The parasitophorous vacuole transmembrane protein, GRA14, can mediate ESCRT-dependent HIV-1 virus-like particle (VLP) budding and internalization of host cytosolic proteins (23). Given the prominent “stalled” intraluminal vesicles within Δ*gra64* tissue cysts, we investigated if host ESCRT components are recruited to the parasitophorous vacuole in a GRA64-dependent manner to facilitate vesicle formation. To that end, we performed a HIV-1 VLP assay as a functional readout to assess GRA64-dependent ESCRT recruitment for HIV-1 VLP release. The HIV-1 Gag p6 domain encoding late domain motifs necessary for ESCRT recruitment was substituted for the GRA64 N-terminal capable of interacting with the host ESCRT machinery (GagGRA64). Deletion of the HIV-1 Gag p6 domain impaired VLP release as previously observed; however, expression of GagGRA64 did not produce VLPs as efficiently as HIV-1 Gag or GagGRA14 (Fig. 6A).

**Figure 6.**
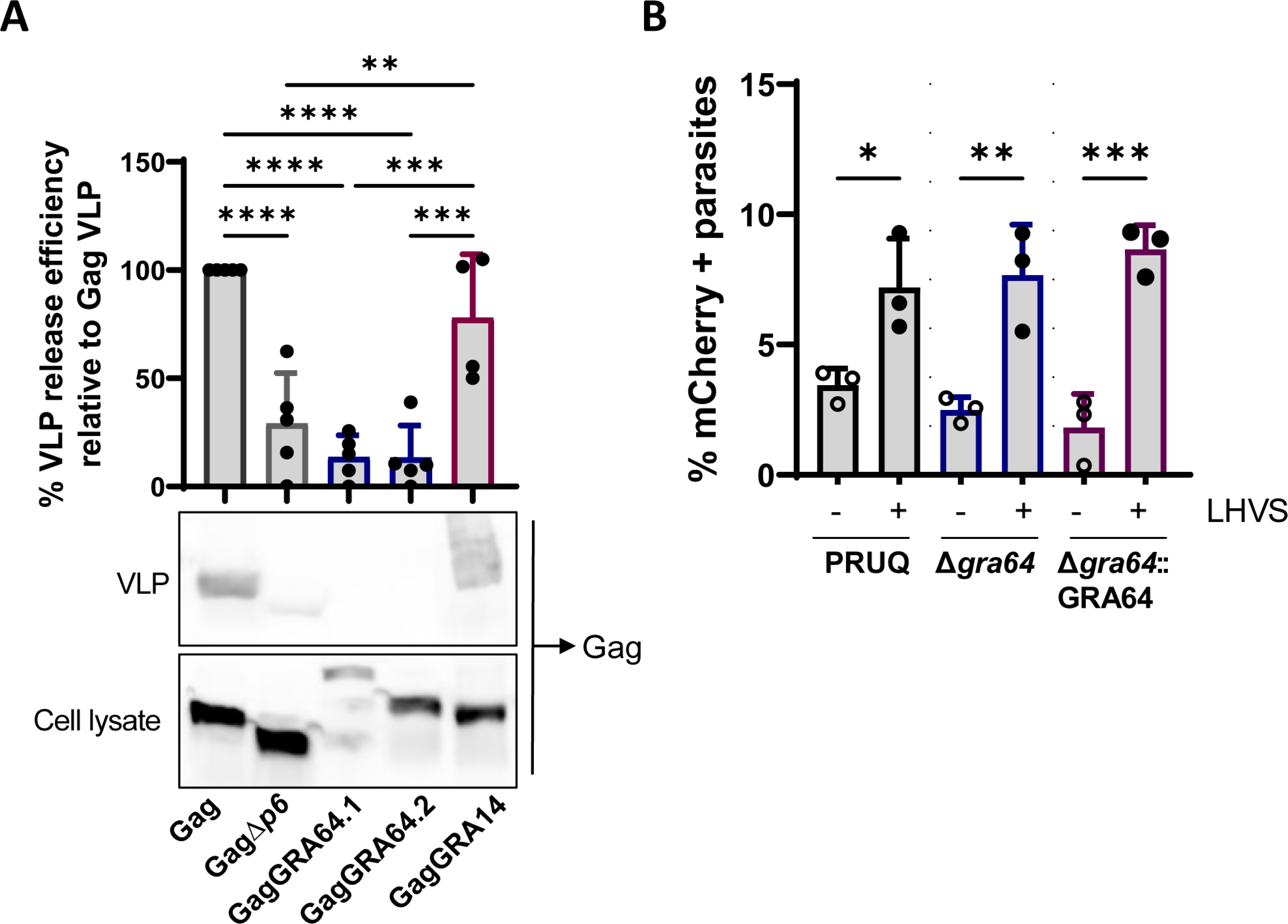
GRA64 does not mediate ESCRT-dependent HIV-1 virus like particle release and is not involved in internalization of host cytosolic proteins. **(A)** Comparison of percent HIV-1 VLP release efficiency between HIV-1 Gag, GagΔ*p6*, GagGRA64 and GagGRA14 relative to Gag VLP. Data represents the mean from n = 4-5 biological replicates. Statistical analysis was determined by a one-way ANOVA test followed by Tukey’s multiple comparison test; **, *P*<0.01; ***, *P*<0.001; ****, *P*<0.0001. **(B)** Quantification of the uptake of host cytosolic proteins in replicating parasites. Analysis of ingestion by PRUQ, PRUQΔ*gra64* and PRUQΔ*gra64*::GRA64 parasites treated with DMSO or LHVS. At least 200 parasites were analysis per sample. Data represents the mean from n = 3 biological replicates. Statistical analysis was determined by a one-way ANOVA test followed by Tukey’s multiple comparison test; *, *P*<0.05; **, *P*<0.01; ***, *P*<0.001.

To further assess the role of GRA64 in vesicular trafficking across the PVM we analyzed internalization of host cytosolic proteins in GRA64-deficient parasites. Inducible mCherry HeLa cells were infected with type II PRU parasites with (PRUQΔ*gra64*::GRA64) or without GRA64(PRUQΔ*gra64*). The infected cells were treated with the cathepsin L inhibitor LHVS prior to harvest at 24 hours post-infection to allow for accumulation of host-derived mCherry within the parasite’s endolysosomal system. The percentage of mCherry-containing parasites was not reduced in Δ*gra64* parasites compared to complemented parasites suggesting that GRA64 alone is not required for the uptake of host cytosolic proteins in replicating parasites (Fig. 6B).

## DISCUSSION

*Toxoplasma* serially secretes effector proteins from the micronemes, rhoptries, and dense granule organelles that aid in invasion, immune suppression, vacuole remodeling, and nutrient acquisition (16). Secreted effectors in both tachyzoite and bradyzoite stages traffic to the PVM or cyst membrane/wall and beyond to the host cell cytosol and nucleus (36, 37). Here, we aimed to identify novel effector proteins using an *in silico* screen to identify gene products with similar properties to known exported effectors and identified a novel dense granule protein (GRA), GRA64. Although GRA64 does not fully translocate across the PVM or cyst membrane and into the host cell as a soluble protein, it does reside at the PVM with the N-terminus exposed to the host cytoplasm. The PVM and cyst membrane are an active parasite/host interface and we observed that GRA64 co-immunoprecipitates with host Endosomal Sorting Complex Required for Transport (ESCRT) components and other parasite GRA proteins during tachyzoite and bradyzoite-staged infection. Proximity-based biotinylation with TurboID also demonstrated ESCRT-proximal proteins within living cells during bradyzoite growth conditions. In addition, GRA64-deficient brain cysts exhibited an abnormal cyst wall structure with enlarged vesicular structures. However, unlike GRA14, GRA64 was unable to substitute the HIV-1 Gag ESCRT-interacting domain to mediate virus-like particle (VLP) release, and GRA64 appears to be dispensable for the internalization of host cytosolic proteins at least under tachyzoite growth conditions (23). However, it is possible that GRA64 recruits the host ESCRT machinery to the parasitophorous vacuole for ESCRT functions at the PVM that remain to be elucidated. This is supported by the fact that the ESCRT accessory protein ALIX is still present at the PVM in GRA14-deficient parasites (23) and the enlarged vesicular structures at the cyst membrane seen in GRA64-deficient parasites. Thus, we speculate that GRA64 is a membrane bound GRA protein that helps orchestrate host ESCRT recruitment indirectly under bradyzoite growth conditions. Co-IP and TurboID experiments confirm interactions between GRA64 and any one of the ESCRT proteins we identified as enriched by these methods however, the limitations of studying these interactions at the host PVM or cyst membrane is due to the lack of functional assays for these interactions.

GRA64 (TGME49_202620) is 330 amino acids in length, possess a signal peptide (1–19), and a transmembrane domain proximal to the C-terminus (268-291) (33). In the hyperLOPIT subcellular proteomics study, GRA64 was suggested to localize to the dense granule organelles (38), a prediction that we confirmed by co-localization with GRA1 (Fig. 1). GRA64 is expressed at all parasite stages but exhibits high expression in tachyzoites and peak expression in bradyzoites (39). It is notable that GRA64 is conserved in *Hammondia*, *Neospora* (two adjacent genes are present), and *Besnoitia*, which are all cyst-forming coccidians and yet have distinct definitive hosts. Our data reveal that GRA64 acts as an integral membrane-bound protein that is expressed at the PVM, IVN (Fig. 1), and the cyst wall (Fig. 2, Fig. 5A). GRA64 is predicted to be intrinsically disordered between the signal peptide and transmembrane domain (Fig. 1B).

Disordered regions are dynamic modules that favor protein-protein interactions, and therefore GRA64 could have multiple roles at the PVM and cyst membrane. GRA64 has verified phosphorylation sites at residues S59, S80, T81, S216 and S218 and a predicted N-glycosylation site at amino acid 55. It is possible that GRA64 activities are further regulated by protein phosphorylation, N-glycosylation and/or another post-translational modification.

The ESCRT machinery is comprised of ESCRT-0, ESCRT-I, ESCRT-II, ESCRT-III, the Vps4 complex, and associated accessory proteins (40), which sequentially interact to drive a wide variety of cellular processes such as membrane repair (41), membrane scission (42), phagophore closure (43), cytokinesis (44), membrane budding (45), and vesicle formation in multivesicular bodies (46). *Toxoplasma gondii* resides within a non-fusogenic parasitophorous vacuole derived from the host plasma membrane (47) and this modified membrane, the PVM, is an active site for vesicular trafficking (48, 49). Since the ESCRT machinery is involved in dynamic membrane processes such as membrane remodeling and scission (50), the ESCRT complex is likely to be involved in cellular processes at the PVM, e.g., PVM repair or host organelle sequestration into the vacuole at the PVM. Recently, ESCRT accessory host proteins (PDCD6IP/Alix, PDCD6, and CC2D1A-a regulator of ESCRT-III CHMP4B) were identified in proximity to the PVM using a targeted PVM domain fused to miniTurbo, a variant of biotin proximity-labeling (51). This finding, and those of GRA14 (23) leave numerous unanswered and exciting questions about the mechanics of ESCRT molecules at the PVM and cyst membrane, and their functional role in the biology of *T. gondii*.

Uptake of host-derived cytosolic proteins occur at the PVM and is partially dependent on the IVN (13), which is a membranous network of tubules that are conduits for moving material such as lipids between the host cell and parasites (52). A recent study demonstrated that several components of host ESCRT-I, ESCRT-III, and ESCRT accessory proteins immunoprecipitated with the vacuole membrane localized GRA14 protein tachyzoite and bradyzoite infected cells (23). Furthermore, a direct connection was discovered between ESCRT and GRA14 at the PVM in the mediation of VLP-budding and in the ingestion of host cytosolic proteins (23). This discovery demonstrates that, in tachyzoites, host ESCRT proteins interact with a parasite effector protein, GRA14 to regulate membrane vesicular budding and the acquisition of host cytosolic cargo. Our findings of GRA64 interactions with host ESCRT-I, ESCRT-III, and accessory ESCRT proteins demonstrate that GRA14 is not the only PVM bound effector that interacts with host ESCRT in the tachyzoite-stage (Fig. 3A). Ongoing and future studies of GRA14 and GRA64 disruption will be essential to determine redundant and/or synergistic roles with host ESCRT in various aspects of nutrient uptake (e.g., recruitment, tethering/binding, ingestion) (53). Furthermore, GRA64 interacts with ESCRT-I, ESCRT-III and ESCRT accessory proteins in the bradyzoite stage in human foreskin fibroblasts (Fig. 3B) and in mouse primary cortical neurons (Fig. 3C). Collectively, these observations demonstrate an emerging paradigm in which parasite proteins interact with host ESCRT at the PVM in tachyzoites and at the cyst membrane in bradyzoites. It is probable that there are other parasite effectors that have yet to be characterized and/or discovered which can also interact with the host ESCRT machinery.

The physical nature of GRA64-ESCRT protein interactions is not yet fully known. We have eliminated a functional GRA64-ESCRT interaction for the VLP budding assay and a dispensable role for GRA64 in the acquisition of host proteins from the cytosol during tachyzoite growth conditions (Fig. 6). These results were not entirely surprising as GRA14, unlike GRA64, possesses motifs that resemble late domain motifs, which recruit ESCRT for HIV budding (54–56). Given the absence of described ESCRT binding motifs in the GRA64 amino acid sequence, it is plausible that recruitment of ESCRT occurs by ubiquitination (57). ESCRT-I can recognize ubiquitinylated proteins that are targeted for degradation in multivesicular bodies (58).

Interestingly TurboID-GRA64 tachyzoites identified a possible GRA64 interaction with the ubiquitin ligase ITCH (Fig. 3D), which is phosphorylated to promote ubiquitination and subsequent degradation of a substrate (59). ITCH can influence lipid metabolism (60), endocytosis (61), viral budding and release (62, 63). Using a Bayesian Discriminant method to predict ubiquitination positions (64), we observed quite a few predicted ubiquitinated lysine residues in GRA64, with the highest ubiquitination score at K227. This theoretical ubiquitin site faces the host cytosol and therefore is a plausible candidate for ESCRT recruitment. Further studies are required to identify the ubiquitin status of GRA64 and whether this modification is required for ESCRT interactions.

*In vivo* cyst defects due to gene knockouts can be hard to characterize as knockouts can result in reduced cyst burdens (65, 66) and increased cyst fragility (67) making cyst purification problematic. However, deletion of GRA64 did not significantly reduce cyst burdens (Fig. S3) and therefore, cysts could be further evaluated for ultrastructural changes (Fig. 5). In the Δ*gra64* cysts, there was an intact cyst wall, a normal IVN, and healthy bradyzoites packed with amylopectin. The knockout of GRA64 did, however, result in the presence of large vesicular structures adjacent to the cyst wall, which we interpret as “stalled-scission events”, in which the inefficient or absent recruitment of ESCRT proteins results in the failure of intraluminal vesicle scission into the mature tissue cysts. The directionality of these large vesicular structures (i.e., whether they are derived from or being targeted to the cyst membrane) and the possible consequences of stalled-scission events are unknown and demand further investigation.

It is not surprising that host ESCRT molecules are intimately involved in the *T. gondii* life stages as the parasite recruits organelles and acquires nutrients from the host such as lysosomes (18), lipids (68) and cholesterol (69). The recent discovery of ESCRT interactions at the PVM and cyst membrane by independent labs corroborate the connection and highlights the complexity of this process. Our findings hint at a possible role for host ESCRT recruitment at mature *in vivo* cyst membranes. This is only the beginning with respect to understanding the relevance of ESCRT recruitment at tachyzoite and bradyzoite vacuolar membranes.

## MATERIALS AND METHODS

### Cell culture

All parasite strains were continuously passaged in human foreskin fibroblasts (HFF:ATCC:CRL-1634; Hs27) in a 37°C, 5% CO_2_ incubator using Dulbecco’s Modified Eagle Media (DMEM, Gibco) supplemented with 10% fetal calf serum, 1% L-glutamine, and 1% penicillin and streptomycin. Cultures were regularly inspected and tested negative for mycoplasma contamination. Bradyzoite induction was performed at the time of invasion by replacing growth media with bradyzoite induction media (50 mM HEPES, pH 8.2, DMEM supplemented with 1% FBS, penicillin and streptomycin) prior to infection of human foreskin fibroblasts with egressed tachyzoites. Bradyzoite induced cultures were maintained in a 37°C incubator without CO_2_, with induction media replaced every 2 days for all experiments.

Mouse primary cortical neurons were harvested from E14 mouse embryos obtained from pregnant C57Bl/6 mice, ordered from Charles River or Jackson Labs. Dissections of E14 cortical neurons were performed as previously described (70). Following dissection, 1x10^6^ - 1x10^7^ cortical neurons were plated onto poly-L-lysine coated 15cm diameter culture dishes and later cultured in Neurobasal Media (Thermo Fisher) supplemented with GlutaMAX Supplement (Thermo Fisher) and B-27 Supplement (Gibco). After 4 days *in vitro* (DIV), cytarabine (ara-C) was added to each culture at a final concentration of 0.2µM to minimize contamination from dividing, non-neuronal cells. Cultures were maintained for up to 16 days by replacing half of the conditioned media with fresh supplemented Neurobasal media every 7 days.

### Cloning and Parasite Transfections

For a full list of primers used for cloning and genetic manipulations, refer to Supplementary Table 1. Briefly, for all Cas9 mediated genetic manipulations, single guide RNAs (sgRNA) targeting the C- or N-terminus of various genes were cloned into the p-HXGPRT-Cas9-GFP plasmid backbone using KLD reactions (New England Biolabs), as previously described (71). 100bp donor oligonucleotides were designed and synthesized (Thermo Fisher) with homologous arms targeting the region of interest and encoding either an epitope tag, stop codons (for knockout transfections), or start codons (for complement transfections) in-frame to the region of interest. Donor sequences for homology mediated recombination with TurboID were generated by PCR using 3xHA-TurboID-NLS_pCDNA3 (kind gift from Alice Ting, Addgene plasmid # 107171) as plasmid template with primers containing overhangs with 40bp homology to the GRA64 region of interest. For epitope tagging GRA64 without Cas9 (ectopic expression), the GRA64 genomic locus was amplified from PruΔ*ku80*Δ*hxgprt* genomic DNA with primers to the C-terminus of GRA64 and 1.5kb upstream of the start codon (the putative promoter region), with overhangs to pLIC-3xHA-DHFR sequences (72). Gibson Assemblies (NEBuilder HiFi DNA Assembly) were subsequently performed to clone PCR amplicons encoding a parasite gene of interest into PCR amplified pLIC-3xHA-DHFR plasmid backbones.

For all transfections, 5x10^6^ - 1x10^7^ PruΔ*ku80*Δ*hxgprt* or RHΔ*ku80*Δ*hxgprt* tachyzoites were electroporated in cytomix (73) after harvesting egressed parasites from human foreskin fibroblast monolayers and filtering through 5µm filters. Selection of transfected parasites was performed with media containing 25µg/mL mycophenolic acid and 50µg/mL xanthine 24 hours post-transfection for 6 days before removing selection media and subcloning by limiting dilution, after sufficient parasite egress was observed. For Cas9 transfections, 7.5µg of uncut Cas9 plasmid and 1.5µg of PCR amplified donor sequence or 280 pmol un-annealed 100bp donor oligos were used per transfection. For ectopic transfections, 10µg of circular plasmid was used for random integration into the genome.

### Immunofluorescence Assays

Human foreskin fibroblast monolayers were grown to confluency on glass coverslips and infected with egressed tachyzoites at an MOI of 1 for most immunofluorescence assays, allowing growth to proceed under tachyzoite or bradyzoite growth conditions (using the media formulation described above). All coverslips were fixed with 4% PFA for 20min at room temperature, permeabilized in a 0.2% Triton X-100, 0.1% glycine solution for 20min at room temperature, rinsed with PBS, and blocked in 1% BSA for either 1 hour at room temperature or at 4°C overnight. For selective permeabilization experiments, coverslips were fixed with 4% PFA for 20min, allowed to cool in PBS at 4C° for 15min, incubated in 0.001% digitonin in PBS for 5min at 4C°, detergent rinsed with PBS, and blocked in 1% BSA for 1 hr at room temperature or overnight. After blocking, coverslips were labeled with antibodies as follows: HA-tagged proteins were detected with rat anti-HA 3F10 (Sigma 1:200-1:500), parasitophorous vacuole with in-house mouse anti-MAG1 (1:500), dense granules with mouse anti-GRA1 (1:1000), and biotin with anti-biotin (Abcam, ab53494, 1:1000). Appropriate secondary antibodies conjugated to Alexa Fluorophores 488, 555, 594, and 633 targeting a given primary antibody species, or streptavidin conjugated to Alexa Fluorophore 488, were used at a dilution of 1:1000 (Thermo Fisher). DAPI counterstain was used to label parasite and host cell nuclei (1:2000). Coverslips were mounted in ProLong Gold Anti-Fade Reagent (Thermo) and imaged using either a Leica SP8 confocal microscope or a Nikon Eclipse widefield fluorescent microscope (Diaphot-300).

### Membrane Fractionation

Human fibroblast monolayers were infected, and parasites were cultured for 2 days under tachyzoite growth conditions. The infected cells were washed and scraped with ice-cold PBS (containing inhibitor cocktail with EDTA, 5 mM NaF, and 2 mM activated Na_3_VO_4_). Infected cultures were lysed by passage through a 27-gauge needle and intact parasites were separated by low-speed centrifugation at 2,500xg for 10min at 4°C. The resulting low-speed pellet was discarded, while the low-speed supernatant (lysate) containing membranous components was separated into soluble and membrane-associated fractions by high-speed centrifugation at 100,000xg for 1.5 hours. The resulting high-speed supernatant (HSS) containing soluble parasitophorous vacuole or cyst components was saved, while the resulting high-speed pellets (HSP) were treated by resuspension in various buffers to free peripheral or integral membrane-associated proteins. Resuspended fractions were centrifuged again at 100,000xg for 1.5 hours to separate liberated proteins in the high-speed supernatant from remaining membrane-bound proteins in the high-speed pellet. All fractions were concentrated by acetone precipitation overnight prior to immunoblotting.

### Immunoblotting

Protein lysates were prepared in radioimmunopreciptation assay (RIPA) buffer from infected fibroblast cultures as specified for each experiment. Laemmli sample buffer was added to all samples and boiled for 5min before loading onto an SDS-PAGE 4-20% pre-cast gradient gel (TGX). Transfer to PVDF membranes (Millipore) was performed in Towbin buffer (20% methanol, Tris/Glycine) for 2 hours at 100V, and blocking in 5% BSA/TBST was performed overnight in 4°C. Membranes were labeled in 5% BSA/TBST with either Streptavidin-HRP (1:10,000, Thermo Fisher), anti-HA peroxidase conjugated antibodies (Sigma, 1:200) or rabbit TgALD1 antibody (1:200, kind gift from Dr. Kentaro Kato) and anti-rabbit HRP antibodies (Thermo Fisher, 1:10000) followed by development of signal with West Pico Plus Chemiluminescent Substrate (Thermo Fisher), or by using LiCor anti-rabbit 680 and LiCor anti-mouse 800 secondary antibodies. Antibodies against TSG101 (Invitrogen Clone 4A10, MA1-23296) and PDCD6 (Proteintech, 12303-1-AP) were also used. Images of labeled blots were collected with a Li-COR instrument (Odyssey Imaging System) or a Bio-Rad Chemidoc Imaging System.

### Immunoelectron microscopy

For immunoelectron microscopy of 3xHA-GRA64 PruQ tagged tachyzoites, samples were prepared from infected human fibroblast monolayers grown under tachyzoite growth conditions for 24 hours. Cultures were fixed in 4% paraformaldehyde and immunolabeled with anti-HA antibody conjugated to both Alexa fluor-488 and a 10nm gold particle using the protocol described above (under Immunofluorescence Assays). After imaging Alexa 488 labeled GRA64, cells were fixed with 2.5% glutaraldehyde and 2.0% paraformaldehyde in 0.1M sodium cacodylate buffer, rinsed with 0.1M sodium cacodylate buffer, postfixed with 1% osmium tetroxide, *en bloc* stained with 2% uranyl acetate, dehydrated in a graded series of ethanol, and infiltrated with LX112 epoxy resin (LADD Research Industries, Burlington VT). Samples were polymerized at 60°C for 60 hours and blocks were popped-off the coverslip. Regions of interest (ROI) were cut out and remounted on a flat BEEM capsule for sectioning. Trimming to the specific ROI was done using Trimtool 45° (Diatome) blocks then serial thin sectioned (70nm) *en face* on a Leica UC7 using a Diatome Ultra 35° knife. Sections were picked up on formvar coated slot grids, stained with uranyl acetate and lead citrate, and photographed using Kodak 4489 film on a JEOL 1200EX TEM.

For electron microscopy of *in vivo* derived tissue cysts, cysts were harvested from homogenized infected mouse brains (4 weeks p.i.) using a Wheaton Potter-Elvehjem Tissue Grinder. Cysts were enriched with 45% Percoll and centrifugation at 26,600xg for 20min at 4°C. The cyst enriched fraction was harvested from the Percoll solution and diluted in PBS prior to a final spin at 130xg for 10min at 4°C. Pellets containing cysts were resuspended in either 2.5% glutaraldehyde and 2% paraformaldehyde in 0.1M sodium cacodylate buffer for morphological analysis or in 4% paraformaldehyde and 0.1% glutaraldehyde in 0.1M sodium cacodylate buffer for immunoelectron microscopy. Samples for morphological analysis were prepared as described above prior to imaging with a JEOL 1400EX transmission electron microscope, whereas samples prepared for immunoelectron microscopy were dehydrated with a graded ethanol series, embedded in Lowicryl HM-20 monostep resin, and polymerized by UV light prior to labeling with anti-HA antibody (Roche, clone 3F10, 1:40 dilution) and 10nm gold bead conjugated goat anti-rat (Electron Microscopy Sciences, 1:100 dilution), or with goat anti-rat only as a negative control. Electron Microscopy images of *in vivo* derived tissue cysts were viewed on a JEOL 1400 Plus using Digital Micrograph software from Gatan.

For tomograms of *in vivo* derived tissue cysts, 250nm thick epoxy sections were picked up onto slot grids and post stained with uranyl acetate and lead citrate. Gold fiducial markers (10nm) were added to the sample prior to imaging to aid in downstream alignment. Tilt series (single axis) were collected on the JEOL 1400 Plus from -60 to +60 using Serial EM software (74). IMOD software was used for alignment and reconstruction of each tomogram as well as the tracing of structures (e.g., vesicles, cyst wall, and parasite membrane) observed in the tomogram to create a model (75).

### Plaque Assays

Parasites were harvested from host cells with a 27G needle and filtered through a 5μm filter to remove host cell debris. Parasite numbers were quantified with a hemocytometer, and 100 parasites from each strain were added in triplicate to wells containing confluent human foreskin fibroblasts in 6-well dishes. Parasites were grown for 14 days before fixing and staining with a 20% methanol-0.5% crystal violet solution. Plaque size was quantified using ImageJ using blinded images and the line tool to separate neighboring plaques. Kruskal-Wallis tests with Dunn’s multiple comparisons were performed to test for significant differences in average plaque size with PRISM 8. Plaque assays were repeated three times with PruQ GRA64 parasite strains, using different batches of parasites and host cells for each independent experiment.

### Co-Immunoprecipitations

For GRA64 Co-IPs from tachyzoite infected cultures, two independent experiments were performed in which 15cm diameter cell culture dishes containing confluent human foreskin fibroblast monolayers were infected at an MOI of 3 with either parasite expressing a 3xHA tag at the endogenous locus of GRA64 (N-terminus), using an equivalent amount of PruQ non-HA tagged parasites each replicate as a control in parallel. Dishes were washed with ice cold PBS 24 hours post-infection and lifted off each dish with a cell scraper in 1mL ice cold lysis buffer (50mM Tris pH 7.4, 200mM NaCl, 1% Triton X-100, and 0.5% CHAPS) supplemented with cOmplete EDTA-free protease inhibitor (Sigma) and phosphatase inhibitors (5mM NaF, 2mM activated Na_3_VO_4_). Scraped cultures were passed through a 27G needle five times and sonicated for 30 seconds total (20% amplitude, 1 second pulses). Sonicated samples were incubated on ice for 30min, supernatant cleared by centrifugation (1000xg, 10min), and incubated overnight in a 4°C rotator with 0.25mg anti-HA magnetic beads (100uL slurry, Thermo Fisher). Following overnight incubation, beads were separated on a magnetic stand and washed twice in lysis buffer and four times in wash buffer (50mM Tris pH 7.4, 300mM NaCl, 0.1% Triton X-100) prior to elution in Laemmli buffer with 50mM DTT, boiling beads for 5min prior to magnetic separation and collection of eluates. Eluates were loaded, washed, and digested into peptides with 1µg of trypsin on S-TRAP micro columns (Protifi) per manufacturer guidelines. S-TRAP peptide eluates were concentrated with a speed vac, desalted in HLB resin (Waters), and concentrated in a speed vac once more prior to LC-MS/MS acquisition.

For GRA64 Co-IPs from bradyzoite and neuron infection conditions, two separate experiments were performed identically to that described above for tachyzoite Co-IPs, except that bradyzoite infections in human fibroblast monolayers were performed with an MOI of 1 in bradyzoite differentiation media, and cultures were maintained for 4 days of infection prior to protein harvesting. For neuron infections, an MOI of 3 and infection period of two days was used prior to protein harvesting, infecting neurons after 14 DIV.

For the reciprocal Co-IP using CHMP4B antibody (Thermo Fisher, rabbit polyclonal, cat no. PA5-64271), 7.5µg of antibody was conjugated to Dynabeads per manufacturer guidelines. Two 15cm dishes were infected with 3xHA-GRA64 tagged PruQ parasites at an MOI of 3, with infection proceeding under tachyzoite growth conditions for 24 hours. Protein was harvested and Co-IPs performed as described above with either CHMP4B-conjugated Dynabeads or unconjugated Dynabeads (with an equivalent weight to that used for CHMP4B antibody conjugation). Eluates were collected and analyzed by immunoblotting, as described above.

### Proximity-Based Biotinylation Protein Preparation

For GRA64-TurboID proximity-based biotinylation experiments, untagged PruQ parasites or parasites expressing TurboID-tagged GRA64 were used to infect confluent human fibroblast monolayers in two 15cm dishes at an MOI of 1 under bradyzoite growth conditions (as described above) for 3 days. Exogenous biotin was supplemented to the media at a final concentration of 150µM. Following infection with biotin supplementation, dishes were rinsed, scraped, and pelleted in PBS (500xg, 10min), after which pellets were solubilized in RIPA buffer supplemented with protease inhibitor cocktail (Roche cOmplete tablets). After 30min of RIPA buffer incubation on ice, insoluble material was cleared from supernatant by centrifugation (16,1000xg for 15min), and supernatant was incubated with streptavidin agarose resin (Thermo Fisher) overnight at 4C° on a rotator. Following incubation, streptavidin resin and bound biotinylated proteins were washed in RIPA urea buffer (50mM Tris-HCl pH 7.5, 8M urea, 150mM NaCl) and subsequently reduced and alkylated with TCEP-HCl and iodoacetamide respectively. On-bead digestion was performed with trypsin and Lys-C proteases, and peptides were harvested from streptavidin resin. Peptides were desalted using C18 tips (Thermo Fisher) prior to liquid chromatography-tandem mass spectrometry acquisition.

### LC-MS/MS Acquisition and Analysis

For peptide samples from all Co-IP and GRA64-TurboID experiments, samples were resuspended in 10 µl of water + 0.1% TFA and loaded onto a Dionex RSLC Ultimate 300 (Thermo Scientific, San Jose, CA, USA), coupled online with an Orbitrap Fusion Lumos (Thermo Scientific). The mass spectrometer was set to acquire spectra in a data-dependent acquisition (DDA) mode. Briefly, the full MS scan was set to 300-1200 *m/z* in the orbitrap with a resolution of 120,000 (at 200 *m/z*) and an AGC target of 5x10e5. MS/MS was performed in the ion trap using the top speed mode (2 secs), an AGC target of 10e4 and an HCD collision energy of 30. Raw files were searched using Proteome Discoverer software (v2.4, Thermo Scientific) using SEQUEST as search engine. We used the SwissProt human or mouse databases (updated January 2020) and the *Toxoplasma* database (Release 44, ME49 proteome obtained from ToxoDB). The search for total proteome included variable modifications of methionine oxidation and N-terminal acetylation, and fixed modification of carbamidomethyl cysteine. Trypsin was specified as the digestive enzyme. Mass tolerance was set to 10 pm for precursor ions and 0.2 Da for product ions. Peptide and protein false discovery rates were set to 1%.

For quantitative analysis, peptide intensity values were log2 transformed, normalized by the average value of each sample and missing values were imputed using a normal distribution 2 standard deviations lower than the mean. Individual peptide fold changes (Tag vs. Control) for a given protein were calculated and averaged to obtain protein fold enrichment. P-values were then obtained from t-distributions and t-values calculated for each protein with at least two detected peptides by treating protein fold enrichment as the sample mean and using log transformed peptide intensity values to calculate the standard deviation, sample size, and degrees of freedom. Data distribution was assumed to be normal, but this was not formally tested. Fold-change and p-value significance cutoffs for both Co-IP and TurboID experiments were arbitrarily selected.

LC-MS/MS data from both Co-IP and TurboID experiments have been deposited onto the public repository Chorus under Project ID 1735.

### Mouse Experiments

Eight-week-old female C57Bl/6 mice (The Jackson Laboratory, Bar Harbor, ME) were infected intraperitoneally with 2000 tachyzoites of a given strain for all acute virulence/survival curve experiments. Mortality was observed daily over 30 days. For cyst burden analysis, brains were collected from C57Bl/6 mice 28-30 days post infection or CBA/J mice (The Jackson Laboratory, Bar Harbor, ME) 28-30 days post-infection. One brain hemisphere or a whole brain from an infected mouse was homogenized with a Wheaton Potter-Elvehjem Tissue Grinder with a 100-150 µm clearance (ThermoFisher) in PBS and an aliquot of the homogenate was viewed under a epifluorescence microscope (Nikon) to count GFP-positive cysts. Kruskal-Wallis tests and Dunn’s multiple comparisons test were performed to test for significance between groups with non-normal distribution with PRISM 8. For groups with normal data distribution, a one-way ANOVA was used to determine statistical significance. A log-rank test was performed to test for statistical significance in Kaplan-Meier survival curves in PRISM 8.

### HIV-1 virus-like particle assay

To generate GagGRA64_Venus, pGag_Venus was linearized using SwaI and SmaI. A fragment encoding the region upstream of the p6 domain from the 664-base pair (bp) to 1456 bp (GagInsert) was generated. Two fragments encoding the GRA64 N-terminus were designed to test the efficiency of GRA64 in substituting the HIV-1 Gag p6 domain. The first fragment encompassed the whole GRA64 N-terminus (GRA64.1) and the second fragment was amplified from 75 bp upstream of the first putative late domain motif encoded in GRA64 and 45 bp downstream of the last putative late domain motif (GRA64.2), the same strategy used to generate the GagGRA14 (23). Fragments were introduced into the linearized Gag_Venus plasmid vector using Gibson Assembly. All plasmids were confirmed by Sanger sequencing.

HIV-1 Gag virus-like particles (VLPs) were collected by ultracentrifugation and analyzed by immunoblot as previously described (23, 76). Lipofectamine 2000 was used to transfect HeLa cells with pRev, pVphu and pGag_Venus constructs. At 18 hours post transfection, the supernatants containing the released VLPs were collected. The samples were filtered and ultracentrifuged at 35,000 rpms for 45min at 4°C to collect the VLP pellets that were further lysed with 0.5% Triton X-lysis buffer. The cell lysates were prepared by lysing the transfected monolayer with the same lysis buffer. The lysates (obtained by loading 100% VLP fraction and 6% cellular fraction on SDS-PAGE gel) were analyzed by immunoblot by probing with human anti-Gag. Band intensity was quantified using Image J. Total Gag corresponds to the sum of cell- and VLP-associated Gag. The VLP release efficiency corresponds to the fraction of Gag that was released as VLP relative to the total Gag. The percentage of VLP release is set to 100% and normalized relative to Gag, which had a % release efficiency of 5.57± 3.6.

### Parasite ingestion assay

Inducible mCherry HeLa cells previously described (23) were seeded in a 6-well plate and induced for 4 days for cytosolic mCherry expression by adding 2 μg/mL doxycycline. The cells were then infected with 1x10^6^ parasites and treated with 5 µM LHVS at four hours post-infection to inhibit the degradation of the ingested material. Parasites were harvested at 24 hours post-infection as previously described (13).

### Ethics Statement

All mouse experiments were conducted according to guidelines from the United States Public Health Service Policy on Humane Care and Use of Laboratory Animals. Animals were maintained in an AAALAC-approved facility, and all protocols were approved by the Institutional Care Committee of the Albert Einstein College of Medicine, Bronx, NY (Animal Protocol 00001451; Animal Welfare Assurance no. A3312-01).

## ACKNOWLEDGEMENTS

We thank members and collaborators of the Weiss lab for their comments, suggestions, and insights on this work. We thank John Boothroyd and Alicja Cygan for useful insights and suggestions on this work. A special thanks goes to the Albert Einstein Analytical Imaging Facility, specifically Xheni Nishku and Timothy Mendez for electron microscopy sample preparation and training, as well as Frank Macaluso for suggestions on electron tomography. We thank the Einstein Laboratory for Macromolecular Analysis and Proteomics for all their assistance with LC-MS/MS preparations and analysis.

This work was supported by P30CA013330, SIG #1S10OD016214-01A1 and SIG#1S10OD023591-01 (Einstein Analytical Imaging Facility), F31AI136401 (J.M.), T32AI070117 (R.B.G), F31AI152297 (Y.R.C), R01AI120607 (V.B.C), R01AI134753 (L.M.W.).

## Author contributions

J.M., R.B.G., and L.M.W. conceived and designed the work, with contributions from Y.R.C., J.R., S.S., V.B.C., and I.C.; J.M. and R.B.G. performed the majority of the experiments, with contributions from V.T. on mouse experiments; T.T. generated the ME49Δ*ku80*Δ*hxgprt* (ME49Q) strain; Y.R.C. performed viral-like particle and parasite ingestion assays; S.S. assisted with proteomic sample preparation and LC-MS/MS analyses; L.G.C. assisted with electron microscopy sample preparations; J.R. and I.C. assisted with electron microscopy and tomogram analyses; J.M., R.B.G. and L.M.W. wrote the paper, with contributions from Y.R.C., S.S. and L.G.C.

**Movie 1. An electron tomogram of ME49Q tissue cyst.** Note the vesicles and tubules in the cyst wall. The dots seen are gold 10nm particles used for alignment during tomogram reconstruction.

**Movie 2. An electron tomogram of ME49QΔ*gra64* tissue cyst.** Note the large vesicles adjacent to the cyst membrane and the lack of smaller vesicles and tubules in the cyst wall. The dots seen are gold 10nm particles used for alignment during tomogram reconstruction.

## SUPPLEMENTAL FIGURES AND TABLES

**Supplementary Table S1.**

List of primers used in this study for CRISPR/Cas9 tagging and cloning of GRA64.

**Figure S1. TurboID-GRA64 LC-MS/MS results from infection of mouse primary cortical neuron cultures.**

TurboID-GRA64 expressing parasites were used to identify proximal proteins in infected mouse primary cortical neuron cultures in two independent experiments. A list of all parasite proteins (blue dots in the volcano plot) with log_2_ fold-enrichment ≥ 2 and a p-value of ≤ 0.1 are listed in the table, while the ESCRT and ESCRT-associated proteins identified as significantly enriched in other datasets (see Fig. 3) are also listed for comparison.

**Figure S2. Δ*gra64* tachyzoites exhibit no growth defects *in vitro* or virulence defects *in vivo*.**

**(A)** Immunoblot demonstrating the absence of GRA64 protein expression in the PruQ knockout strain (Δ*gra64*) and comparable amounts of protein expressed in the PruQ parental (3xHA-GRA64) and complemented PruQ strain (GRA64-COMP). TgALD1 was used as a parasite specific loading control.

**(B)** Plaque assays in human fibroblasts monolayers cultures following 14 days of infection under tachyzoite growth conditions demonstrate no statistically significant difference in parasite growth, as measured by plaque sizes between each strain in three independent experiments. Representative images of plaques are provided for each strain beside the violin plots.

**(C)** C57Bl/6 mouse mortality was recorded over a span of 30 days post-intraperitoneal infection, using 2000 tachyzoites of each strain to infect 10 mice each. No significant differences in the survival curves were noted in this experiment (n.s.).

**Figure S3. Disruption of GRA64 does not significantly affect tissue cyst burdens.**

**(A-B)** Cyst burdens were measured from the brains of chronically infected CBA/J mice (30 days post-infection) using (A) PruQ strain or (B) ME49Q strain parasites. (A) *P*-values were calculated from a One-Way ANOVA test for significance in Replicate 1 and a Kruskall-Wallis test in Replicate 2 (as data were not normally distributed in Replicate 2). The data indicate a trend of fewer cysts formed during Δ*gra64* infection compared to parental and complement strains in both experiments. No significant differences in cyst burden between parental and complement strains are present in either experiment. (B) *P*-values were calculated from a one-way ANOVA test for significance in both replicates. The data shows higher cyst numbers using a more cystogenic strain; however, no significant differences in cysts formed during Δ*gra64* infection was noted compared to wild-type, parental and complement strains in both experiments.

**Figure S4. Disruption of GRA64 *in vivo* brain cyst exhibits large vesicular structures that have close contact with the cyst membrane.**

Cysts were purified from the brains of chronically infected CBA/J mice (30 days post-infection) using the (A-B) ME49Q and (C-D) ME49QΔ*gra64* parasites, prepared for EM serial imaging, and reconstructed. (A and C) The vesicles (shown in yellow, blue, green, and cyan), cyst wall (shown in magenta), and parasite membrane (shown in red) were traced using 3dmod within the IMOD software from the cyst shown in Movie 1 and 2. (B and D) A reconstruction model is shown comprised of each serial image traced for the ME49Q and ME49QΔ*gra64* cysts.

## Supplementary Dataset 1. LC-MS/MS Data from Co-IP experiments

Data are present in eleven different tabs. The “Summary” tab for tachyzoite, bradyzoite, and neuron lists the calculated protein fold change (GRA64-3xHA/Control) and -log_2_ p-values for all the proteins detected in each replicate. The “Replicate 1/2 Analysis” tab for tachyzoite, bradyzoite, and neuron demonstrates data transformation steps and equations used to determine average protein fold change and -log_2_ p-values. The “Replicate 1/2 Raw Data” tab provides information on search engine identification quality parameters. This dataset has been deposited into the mass spectrometry open access repository Chorus under Project ID 1735.

## Supplementary Dataset 2. LC-MS/MS Data from TurboID experiments

Data are present in seven different tabs. The “Results Summary” tab for human foreskin fibroblast and neuron lists the calculated protein fold change (TurboID-GRA64/Control) and - log_2_ p-values for all the proteins detected in each replicate. The “Replicate 1/2 Analysis” tab for human foreskin fibroblast and neuron demonstrates data transformation steps and equations used to determine average protein fold change and -log_2_ p-values. The “Raw Data” tab provides information on search engine identification quality parameters for replicate 1 and 2. This dataset has been deposited into the mass spectrometry open access repository Chorus under Project ID 1735.

